# Tactile responses in the mouse perirhinal cortex show invariance to physical features of the stimulus

**DOI:** 10.1101/2025.08.15.670508

**Authors:** Rejwan Hamza Farhad Salih, Luis Fernando Cobar, Francois Philippe Pauzin, Davide Zoccolan, Maximiliano Jose Nigro

## Abstract

Sensory processing in the cortex follows a hierarchical structure, transforming fine-grained stimulus features into abstract, invariant representations. These transformations have typically been characterised in visual pathways ranging from primary visual cortex to inferotemporal cortex. Less is known about how stimulus representations are formatted in other modalities and downstream regions such as the perirhinal cortex (PER), which receives processed sensory input from all modalities and is involved in object recognition. To address this, we investigated neural activity along the somatosensory hierarchy in awake, head-fixed mice receiving tactile whisker stimulation on either side of the face. PCA and stimulus-baseline decoding analyses revealed that PER activity was similar across stimulated sides, while this was not the case in primary somatosensory cortex (wS1). We also performed decoding analyses for the side of stimulation and the stimulus’ movement direction and found that wS1 and secondary somatosensory cortex could reliably decode both features. In contrast, PER was shown to be invariant to both features. We also clustered neurons based on their response dynamics, which indicated that the majority of PER neurons did not encode the basic stimulus features. These findings demonstrate that PER is a critical node in the somatosensory processing hierarchy, encoding tactile information in an invariant and abstract format. Together, these insights contribute to a more refined understanding of how perceptual processing in PER contributes to the various cognitive processes it is utilised for.

**Significance Statement:** Sensory information is processed in stages that transform detailed signals into complex, stable representations. The perirhinal cortex (PER) is at the apex of these stages and is involved in recognizing objects, a cognitive process that requires both perceptual and memory-based processing. While PER’s role in the latter is established, evidence for the former is conflicting. To investigate this, we recorded PER activity in mice passively experiencing simple tactile stimulation with two pairs of feature combinations. Along the cortical somatosensory hierarchy invariance of stimulus representation emerged in PER, while individual features were represented in earlier cortices with high fidelity. This suggests that PER contributes to the formation of invariant representations of sensory stimuli that is a foundation for object recognition.

## Introduction

Sensory information is sequentially processed along a hierarchically organized network (Felleman and Van Essen, 1991). This organisation enables the representation of objects to emerge in higher order cortices from initially fine-grained stimulus features represented in primary sensory areas. In the visual system, this corresponds to the responses of simple cells in primary visual cortex, which represent stimulus edges, colour, contrast, etc (Hubel and Wiesel, 1962; Movshon et al., 1978; Albrecht and Hamilton, 1982; Skottun et al., 1991; Friedman et al., 2003; Li et al., 2006). Along the following processing stages, neurons encode more complex combinations of visual features and are invariant to increasingly larger transformations of visual stimuli. For example, the IT cortex encodes objects in a manner that is invariant to object rotation, translation, scaling, and so forth (DiCarlo, Zoccolan, et al., 2012)), suggesting that hierarchical processing is a key component of visual object recognition (DiCarlo and Cox, 2007).

Hierarchical processing is not unique to visual information, having been observed across sensory modalities (Hsiao et al., 2002; Asahina et al., 2009; Sharpee et al., 2011; Zoccolan, 2015; Brandao et al., 2021; Schiltz et al., 2024). In the somatosensory modality, neurons in layer 4 of the vibrissal primary sensory cortex (wS1) show narrow receptive fields that encode the identity of single whiskers (Voelcker et al., 2022). Receptive fields become broader in layer 2 where neurons show multiwhisker responses and whisker-invariant encoding of force deflection (Voelcker et al., 2022). Along the somatosensory cortical hierarchy, categorical responses emerge in the secondary somatosensory cortex (S2) where broader receptive fields might contribute to the coding of stimulus identity or task-dependent behavioural features (Helmchen et al., 2018; Rossi-Pool et al., 2021).

The perirhinal cortex (PER) is a cortical region within the medial temporal lobe (MTL) that receives uni- and multisensory inputs from all sensory modalities (Kealy and Commins, 2011). Early investigations in macaques and rats demonstrated its role in visual object recognition (E Murray and Mishkin, 1986; Zola-Morgan et al., 1989; Meunier et al., 1993; Eacott, Gaffan, et al., 1994). The perceptual role of PER in object recognition has been explored in several macaque and rat lesions studies, suggesting an important role of this region in supporting the discrimination of configural stimuli (i.e., visual patterns or 3d objects that can only be distinguished by taking into account multiple features) (M Buckley and Gaffan, 1998; M J Buckley et al., 2001; Eacott, Machin, et al., 2001; T J Bussey et al., 2002; T J Bussey et al., 2003; Bartko et al., 2007b; L M Saksida et al., 2007; Ramos, 2014). However, neurophysiological recordings in freely moving rats have provided contrasting results on object representation in PER (Burke et al., 2012; Fiorilli, Marchesi, et al., 2024). Finally, while some PER neurons respond to visual objects in a way that is more consistent with mnemonic functions (i.e., by encoding object response category), others display a graded tuning for the objects’ perceptual features (Ahn and I Lee, 2017). These studies demonstrate how hard it is to disentangle perceptual from mnemonic contributions especially in freely moving animals engaged in complex tasks as compared to a classical sensory physiology experimental setting.

We performed neurophysiological recordings along the somatosensory cortical hierarchy of awake head-fixed mice receiving tactile stimulation on either side of the face by moving an object in the whisker pad. This paradigm allows to test for invariant responses of two well-controlled stimulus features i.e., side of the face and position of the object in the rostro-caudal axis. We observed that responses of PER neurons were more similar between the four stimulus features than in wS1. Decoding approaches demonstrated that the population of PER neurons show invariant responses to the two physical features presented. These invariant responses emerge in PER, suggesting that local processing underlies this transformation. Our results provide insights on the mechanisms by which transformation of sensory representations in PER might contribute to building abstract representations of stimuli such as the presence of objects, context or task structure (D G Lee et al., 2024).

## Materials and Methods

### Subjects

Data were collected from *n* = 9 experimentally naive C57BL/6 mice, four males and five females between 12-24 weeks of age at the time of head-bar implantation surgery. Before surgery, all mice were housed with up to four littermates. After surgery, each mouse was individually housed in a large plexiglass cage. All mice were kept on a 12-h light–12-h dark schedule, with strict control of humidity and temperature. All procedures were performed in accordance with the Norwegian Animal Welfare Act and the European Convention for the Protection of Vertebrate Animals used for Experimental and Other Scientific Purposes. Protocols were approved by the Norwegian Food Safety Authority (FOTS ID 25405, 31049).

### Surgery

Anaesthesia was induced with isoflurane (4%, maintained at 1.5% but minor adjustments were made during the surgery), followed by subcutaneous injections of Buprenorphine and Meloxicam as well as a subcutaneous injection of Bupivacaine at the incision site prior to the incision. Once the skull was exposed, fiducial marks were made on the skull at AP −2.0 mm and ML +3.6 mm on the right hemisphere for the probe targeting the Perirhinal (A35 and A36) cortex and at AP −1.4 mm and ML +3.00 on the right hemisphere for the probe targeting wS1. Afterwards, a plastic ring cut from pipette tips (∼2.5 mm diameter) was attached to skull using Vetbond and once it was firmly attached, it was covered with Kwik-Cast, to protect the skull, but to leave the fiduciary marks clear for later access. Afterwards, a ground screw, a screw with a gold pin (WPI Pin 5482) soldered to the screw’s head, was affixed to the skull at AP −2.0 mm and ML +2.50 mm on the left hemisphere. For additional support, two extra screws were attached to skull and a stainless-steel head bar was attached to the skull using Optibond. After the headbar was attached, the rest of the exposed skull was also covered with Optibond. After acclimatization in the head-fixation apparatus, which usually lasted about 1.5 – 2 weeks, the animals were again anaesthetized with isoflurane and placed on the stereotaxic frame. The Kwik-Cast and pipette ring were removed, exposing the fiduciary marks for both wS1 and PER targets. Afterwards, using a drill bit, the bone was made as thin as possible, and once it was very thin, the bone was gently broken using a blunt glass pipette, thus completing the craniotomy. After both craniotomies were completed, they were both covered with Kwik-Sil, and the mouse was allowed to recover for approximately 1 hour. Afterwards, the mouse was placed on the head-fixation apparatus, the ground pin was connected to the grounding circuit from the recording setup and the craniotomies were exposed. Each Neuropixel 1.0 probe (Phase 3B) was inserted into the brain at approximately 10 µm per second until it reached the planned depth (DV −3.00 mm for PER probe and −2.16 mm for the wS1 probe). The coordinates of the PER probe target was the rostral PER, which has been shown to be a projection target of wS1 (Aronoff et al., 2010). Once the probes were in their final location, the craniotomies were filled with 0.9% Saline solution to avoid drying out the tissue. The probes were inserted into the brain and kept there for 30 minutes before starting the recording, to minimize drifting artifacts and improve signal stability. The experiment lasted for about 55 minutes for 4 mice, and about 80 minutes for the rest of the mice. Afterwards, mice were euthanised after a single recording session.

### Stimulation protocol

Tactile stimulation was delivered using a pair of glass pipettes (Harvard Apparatus, 30-004 Glass Capillaries, GC120F-10) attached to two piezo actuators (THOR Labs PB4NB2S) under the control of a piezo controller (MDT693B). The glass pipettes were bended at 90° in the extremity towards the mice and positioned approximately ∼4 mm away from the face of the mouse and were in a position orthogonal from the whisker natural resting orientation. Each trial featured a pseudorandom choice of the side being stimulated, defined as ipsilateral or contralateral to the recorded hemisphere. Four of the nine mice were also administered trials with bilateral stimulation, but those trials were not included for analysis. Stimulation times were recorded using custom-made Simulink scripts and aligned to electrophysiological recordings offline.

Stimulations began with a rostro-caudal movement (a displacement of ∼ 0.59 mm, measured at the tip of the pipette closest to the mouse whiskers), followed by a 500ms pause, and then a caudo-rostral movement back to the starting position for another 500ms (each movement from the pipette occurred within 5ms). This 1Hz stimulation cycle was repeated another four times, resulting in a 5s stimulation window. This train of stimulations was followed by a 15s (N=4) or 5s (N=5) baseline period where the pipettes remained stationary. The difference in baseline periods was made to increase the number of stimulation periods within the experimental session. In each session, there were a total of 100 and 400 trials, respectively, split near-evenly between ipsi- and contralateral. For experiments with 100 trials, contralateral stimulations were 47.25 ± 4.80 and ipsilateral stimulations were 48.50 ± 4.15; for experiments with 400 trials, contralateral stimulations were 202 ± 4.73 while ipsilateral stimulations were 197.6 ± 5.49. Each session also featured a control block in which the stimulation protocol was repeated but with the pipettes moved away from the whiskers. In this block there were 5-7 trials for the former cohort, and around 80 for the latter cohort. Analysis of the control period is not included in this manuscript.

### Histology

After the experiment was concluded the probe was removed and the craniotomies were covered using Kwik-Sil and the mouse was euthanized with an injection of pentobarbital perfused transcardially with 0.9% Saline solution followed by 4% formaldehyde (v/v) diluted in PBS. The brains were extracted and stored in 4% formaldehyde in PBS for 24 hours. Afterwards, the brains were transferred to 30% sucrose and stored for at least 48 hours. The brains were then frozen and cut into 35 µm coronal sections with a freezing microtome (Thermo Scientific Microm HM430) and imaged with a widefield fluorescence microscope (AxioScan Z1, Zeiss).

After the images were acquired, we identified the sections where the probe entered the cortex and where the tip of the probe was located. We then used open-source software HERBS (Fuglstad et al., 2023) to estimate the anatomical location of each channel and identify which channels were in PER and wS1. Subsequent analysis only used units recorded only on channels located on these two cortical regions.

### Neural recordings

Extracellular recordings were made by acutely inserting a silicon probe into each craniotomy, one targeting the barrel cortex and the other one targeting the perirhinal cortex. Probe insertions were performed while the animal was head-fixed in the stimulation setup.

All recordings were made with Neuropixels 1.0 probes (Phase 3B)(Jun et al., 2017). Raw voltage traces were filtered, amplified, multiplexed, and digitized on-probe and recorded using SpikeGLX (https://billkarsh.github.io/ SpikeGLX). Voltage traces were filtered between 300 Hz and 10 kHz and sampled at 30 kHz with gain = 500 (AP band) or filtered between 0.5 Hz and 1 kHz and sampled at 2.5 kHz with gain = 250 (LFP band). All recordings were made using the channels closest to the tip of the probe (Bank 0). The probe’s ground and reference pads were shorted together and soldered to a gold pin, which was then connected to a gold pin that had been soldered to a skull screw on the animal. All recordings made were in external reference mode, using the skull screw as external reference.

The probe targeting PER was inserted at 90 degrees, while the one targeting wS1 was at 75 degrees from the horizontal plane. During the recordings, the mouse was free to run or to stay immobile on the running disk. The timing of the movement of the piezo actuators was recorded using custom-made Simulink scripts and an Arduino UNO. For synchronization of neurophysiological data and other behavioural and experimental variables, an additional Arduino UNO was used to generate digital pulses, which were sent to the Neuropixels acquisition system (via direct TTL input) and the other Arduino UNO. After recording, all data were aligned off-line using custom-made MATLAB scripts.

### Unit classification and inclusion criteria

After spike sorting using Kilosort 2.5 and Phy (Rossant, Kadir, et al., 2016; Rossant, Harris, et al., 2021; Pachitariu et al., 2024), in which each unit’s various features, including, ISI violations, similarity scores, percentage of missing spikes, stability during the recording session and abnormal waveform were assessed in order to merge or separate different clusters. Once manual curation was completed the resulting neurons were required to pass three criteria to be included for further analysis: i) ≥ 500 spikes in total, ii) Presence ratio, defined as the fraction of 60s bins tiling the whole recording duration containing at least one spike, ≥ 0.9, iii) coefficient of variation of firing rates calculated using 20s bins ≤ 1 (Buccino et al., 2020). Additionally, the units were split into regular- (RS) and fast-spiking (FS) using their waveform duration as the guiding metric. Namely, RS and FS units had a duration ≥ 0.4ms and *<* 0.4ms, respectively. For all the analyses in this work, we used the entire population of both RS and FS units together.

### Single-unit activity

Mean activity traces were constructed using a bin width of 1ms and averaging spike counts over all trials (ipsi- or contralateral). These binned firing rates were then smoothed using, unless otherwise stated, a gaussian kernel with a width of 20ms.

### Population representation of the stimulus sequence and PCA analysis

To understand how PER neurons encoded the tactile stimulus sequence, we mapped each time bin of the stimulation epoch on the representational space defined by the activity of the recorded neurons. More specifically, the firing rates of each neuron were trial-averaged (per trial type; ipsi- or contralateral) over a window of [−2.5, +7.5]s relative to stimulus onset, using bins of 1ms width. These mean firing rates were then smoothed using a gaussian kernel of 20ms width. To minimise any bias from high firing rate units, the smoothed firing rates were centred and scaled through z-scoring. This also served to prevent baseline firing differences to artificially separate the dynamics in state space. These smoothed and z-scored firing rates were combined for each recorded region, producing two 2D matrices, one for each trial type. Each matrix was *n* × *T* in size, where *n* is the number of recorded neurons in a given region, and *T* is the number of 1ms time bins (10000) along the [−2.5, +7.5]s window. Ultimately, in each matrix, a column was a population vector representing the activity of the recorded PER population at a specific time over the course of the stimulation epoch.

The dimensionality of this population representation was then reduced by applying PCA. For each region, the two matrices were analysed in two ways (Figure 3A). In the first approach, we wanted to determine whether neurons contributed similarly to the population dynamics produced in each trial type. To do so, we applied PCA separately across the rows of the two matrices, extracting, in each case, the first principal component (PC) only. This reduced each matrix to a size of 1 × *T* and provided a weight array of size *n*. These weights indicated how much each neuron contributed to the first PC. For each neuron, this resulted in a pair of weights, one from the first PC of each matrix. Using these weight pairs across neurons, we applied linear regression to determine how correlated they were.

In the other approach, our aim was to visualize and compare the population dynamics for each trial type over the course of stimulus presentation. This required mapping the population activity recorded in each time bin on the same representational space. Thus, the two matrices were concatenated time-wise and then PCA was applied to this combined matrix of size *n* × 2*T* . We then retained the first three PCs to reduce the dimensionality of the representational space in each trial type (the two matrices) from a size of *n* × *T* to 3 × *T* . This allowed plotting the population dynamics in a 3D PC space that was common to the two trial types.

### K-means clustering of neural response patterns

The K-means methodology used in this work was adapted from a previous study (Adler et al., 2012). As done for the analysis described in the previous section, also in this case, we first built a population representation of the stimulus sequence. First, smoothed average firing rates were computed over a window of [−1, +6]s relative to stimulus onset (per trial type). Compared to the approach described in the previous section, here we included less of the baseline periods. The rationale was to ensure that clusters would be derived primarily on the basis of firing activity within the stimulation window. For each neuron, these firing rates were subtracted by a baseline firing rate that was computed over a 3s window, starting prior to stimulus onset. These baseline-subtracted, smoothed firing rates were then z-scored according to the mean and standard deviation of each neuron. This yielded a temporal sequence of population vectors that, as described in the previous section, were organized in a *n* × *T* in size, where *n* is the number of neurons of a given region, and *T* is the number of 1ms bins (7000, in this case). Prior to K-means application, the temporal dimension was reduced to three dimensions by applying PCA along the columns of the matrices. Accordingly, this produced a final output of size *n* × 6 as it was applied to each stimulus type separately.

It is worth highlighting the different application of PCA here, as compared to the analysis described in the previous section, where the dimensionality reduction was performed across the rows of the matrices containing the population vectors. In that case, this allowed projecting the trajectory of the population activity, during the course of stimulus presentation, over a lower dimensional neuronal representational space. Applying the PCA across the columns of the population vectors’ matrixes allows instead projecting each neuron over a low dimensional space defined by the combinations of stimulus epochs that give rise to each principal component. This, in turns, enables identifying subsets of units that are similarly activated by specific temporal features of the stimulus sequence by applying a clustering analysis.

To this end, the output matrices produced by the PCA were clustered using K-means, starting with *K* = 2 clusters up to *K* = 10. For each value of *K*, the output underwent a silhouette analysis (Rousseeuw, 1987). This produces a value (silhouette score) between 1 and −1 for each neuron based on how strongly the neuron is within its assigned cluster and how separated it is from neurons of other clusters. We compared the spread of silhouette scores both within and between clusters. These comparisons were done with reference to the mean silhouette score. We also produced an elbow plot, which shows a monotonic decrease in the sum of squared distances of each neuron to its cluster center with increasing *K*. The “elbow” of this plot can be used to estimate *K*. These two approaches were applied, along with visual inspection of the resulting clusters, to determine the appropriate value of *K*. For PER, this value was determined to be *K* = 4 clusters.

### Decoding methodologies

All decoding analyses were performed using support vector machines (SVMs) through the scikit-learn function SVC (Pedregosa et al., 2011). For all implementations, we set the regularization parameter to *C* = 0.0001 and used a linear kernel to produce linear separating hyperplanes. Unless otherwise stated, all reported classification accuracies are obtained by averaging performance across 100 independent runs, where each iteration includes a resampling of neurons and randomised partitioning of trials. Also, firing rates were binned using 5ms windows instead of 1ms and no smoothing was performed.

### Stimulus-baseline decoding

Here, we used SVMs to determine whether population responses recorded during the stimulus epoch were distinguishable from those associated to baseline activity (Figure 4A; “within-conditional”). For each region and stimulus type, we randomly selected *N* = 30 neurons and split trials into two groups. Subsequently, the data in each group were trial-averaged, producing two 2D matrices of 5ms-binned firing rates. Individual SVMs were trained and tested across time using a time window of 150ms with a step size of 50ms. Starting at stimulus onset, thirty 5ms bins of one group were used as training data, with the corresponding data of the other group used for testing (i.e., to measure the performance of the classifiers). In addition to stimulus window data, thirty adjacent 5ms bins were chosen from the baseline window starting at a random point within the window. This choice was the same for both groups and provided training and testing samples for classifying baseline data. For both the training and testing data, the corresponding class labels were thirty 1s and thirty 0s, for stimulus and baseline window data, respectively. Performance was calculated as the percentage of testing data that was correctly classified by the SVM. This procedure was repeated but with the training group now forming the testing group, and vice versa, thus implementing a two-fold cross-validation measurement of classification performance. The whole procedure was repeated across the entire stimulus window. Since the resulting accuracies are specific to the given sample of 30 neurons and choice of trials in each group, we repeated this entire procedure 100 times and averaged the decoding accuracies obtained across such 100 runs in each time window.

To evaluate the statistical significance of these final accuracies, we used permutation testing to generate an empirical null distribution. To do this, we used the same 100 samples of neurons and trial splits to ensure robustness. First, a base class label vector was randomly shuffled 1000 times. This vector contained thirty 1s and thirty 0s, in accordance with the decoding data sampling procedure. In each shuffling iteration, the resulting label vector was used to train the decoders, but the unshuffled label vector was used for testing. An average accuracy was calculated by evaluating performance over all 100 runs of the decoding. Through this procedure, we generated 1000 null averaged accuracies in each time window. In each time window, the distribution of these null averaged accuracies can be used to generate confidence intervals to determine if the corresponding unshuffled average accuracy is significant. However, since we would therefore test for significance across multiple time windows, we performed multiple comparisons correction using the maxT approach (Westfall and Young, 1993), which controls the family-wise error rate.

To illustrate this approach, consider that we obtain a 1000 × *t* matrix containing null averaged accuracies, where *t* is the total number of 150ms time windows within the stimulation window. For each time window, we compute the average over the 1000 accuracy values, resulting in a vector, *V* , of mean null accuracies of length *t*. Next, for each shuffle and time window, we calculate the deviation from the corresponding time window’s mean null accuracy, yielding a deviation matrix of size 1000 × *t*. For each row (i.e. shuffle permutation), we extract the maximum deviation values across the *t* time windows, producing a vector of length 1000. These maximal deviations reflect the most extreme fluctuations expected under the null hypothesis and are used to construct a null distribution of extrema. To compute corrected 95% confidence intervals, we compute the 95th percentile of the maximum deviation distribution. This value is then added to *V* to yield upper confidence bounds. An unshuffled average accuracy is deemed significant if it exceeds the corresponding corrected bound in a given time window.

### Cross-conditional stimulus-baseline decoding

This approach was similar to the stimulus-baseline decoding approach described above (Figure 4A; “cross-conditional”). As before, each decoding iteration involved a random selection of *N* = 30 units and trial partitions, which, again, was repeated 100 times. Across trial types, this results in two sets of two 2D matrices. Performance, and statistical significance, was evaluated in the same manner as above, but here, for each classification run, we also evaluated performance for cases where the training group was from trials of one type (e.g. ipsilateral) but the test group used trials of the other type (e.g. contralateral). We describe these cases as “cross-conditional”, whose performance is compared to the within-condition approach (i.e., training and testing on the same trial type, as described above).

For each run (i.e., each selection of 30 units and trial partition), these cross-conditional and within-condition classification accuracies were compared by subtracting the latter from the former. The resulting differences were then averaged and their standard deviation was computed across time. We also displayed the distribution of these differences as a kernel density estimate plot. We also calculated the average value of these differences across time points and determined whether it was statistically significant by performing a two-tailed paired permutation test. Specifically, for the two curves that were subtracted to create the distribution of differences, we created a surrogate pair of curves. To do this, in each time point we randomly chose whether to exchange the values in the two curves or keep them the same. This procedure was repeated 1000 times, producing 1000 pairs of surrogate curves. The difference between pairs was calculated and then averaged over time to produce 1000 values of average accuracy differences. To centre the resulting distribution, we subtracted its mean from each value. This mean was also used to centre the real average difference. Then, a two-sided p-value was calculated by counting the number of samples whose absolute centred values were greater than the absolute value of the centred real average difference. We divided this count by the number of samples (1000) after adding 1 to both values for continuity correction.

### Decoding of stimulus type

This approach was also similar to the stimulus-baseline decoding approach but the aim was to measure whether the neuronal representation in a given area was able to support the discrimination of stimulus type (i.e., ipsilateral vs. contralateral). Each decoding iteration involved a randomly selected *N* = 30 units and split trials into two groups for each stimulus type, as before. Here, one of the two groups from each stimulus type were used as training data, creating 30 × 2 data samples for training in each 150ms window. The remaining group from each stimulus type constituted the testing data. All other steps were identical. For testing statistical significance, the permutation test was the same as described above for the individual decoding accuracy curves.

### Decoding of stimulus direction

To decode the stimulus direction, we split the entire 5s stimulation window into ten 500ms windows. Each window begins relative to the movement of the stimulating pipette. Given the square-wave pattern of motion (see Figure 1A), five of these windows correspond to a rostro-caudal direction of motion, with the remaining five to a caudo-rostral direction.

**Figure 1:**
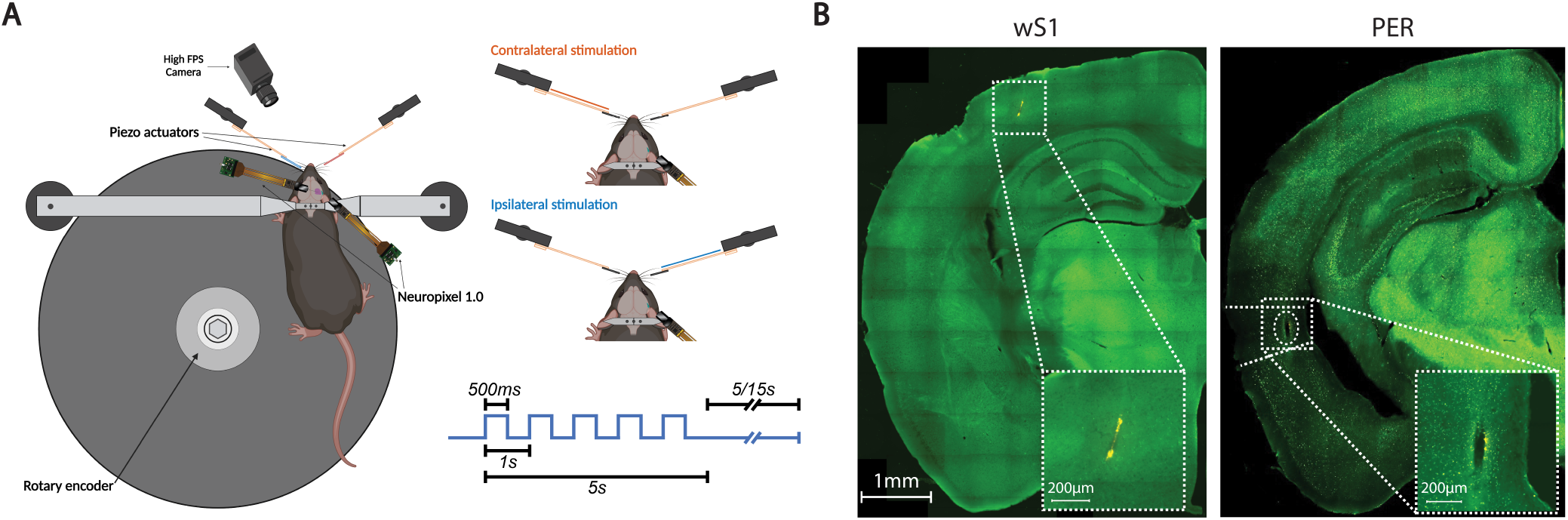
Overview of the experimental setup. **A:** Schematic of the experimental apparatus and stimulation protocol. Mice were head-fixed on a running disc with a rotary encoder that measured their speed along with a camera placed in front of them. The top-right panel illustrates the labelling of trial types with the bottom-right explicating the pipette movement during the stimulation window. **B:** A representative example of probe trajectory for each targeted region, wS1 on the left and PER on the right.

In this case, to train and test the classifiers we used a shorter time window of 30ms (rather than 150ms) with a step size of 10ms. Instead of sweeping across the entire stimulation window, our sampling was relative to the start of each of the 10 movements in the stimulation window. For example, for the first decoding window, we sampled the first 30ms of data after each of the movements, which provided 10 sets of 6 firing rate vectors. The firing rate vectors were generated as done for the previous decoding analyses described above. Since movement direction alternated, there were 5 sets of data for each of the two movement directions, i.e., 30 firing rate vectors each. For class labels, we used two vectors of size 30, one containing 0s for the data from one movement direction, and the other containing 1s for the other movement direction. The calculation of decoding accuracies was identical to the above, namely that two-fold average accuracy was averaged over 100 decoding iterations for each time window. The shuffling procedure was also identical.

### Statistical analysis

Strength of association for linear regression was evaluated using the coefficient of determination (*R*^2^), with the associated *p*-value determined using the Wald Test. For decoding analyses, statistical significance was assessed using non-parametric permutation tests. Correction for multiple comparisons across time windows was implemented using the maxT method (Westfall and Young, 1993), which controls the family-wise error rate. Significance thresholds were derived from the 95th percentile of these maxT-corrected empirical distributions. For comparisons of the cross- and within-condition decoding accuracies, *p*-values were derived from the corresponding empirical distributions, with continuity corrections applied to avoid zero *p*-values. Unless otherwise stated, results are reported as mean values. All error bars indicate the standard deviation, except for the K-means cluster-averaged response traces, which instead indicate the standard error of the mean.

### Code accessibility

All code used in this manuscript can be accessed in the GitHub repository (https://github.com/rejwansalihntnu/ peri_tactile)

## Results

### Experiment structure and single-unit isolation

The experimental paradigm featured head-fixed mice (N=9) on a running disc receiving tactile stimulation through movements of glass pipettes placed near each side of the face inside the whisker pads (Figure 1A). In each trial, one of the two pipettes moved in a square-wave pattern at 1Hz for 5s. Each cycle featured a rostro-caudal movement, followed by a 500ms pause before returning to the original position for another 500ms. Each trial ended with a baseline period of 5s (N=5) or 15s (N=4).

Neural recordings were made by implanting two neuropixels 1.0 probes per mouse. One probe targeted wS1 and the other targeted PER (Figure 1B). Over the 9 mice, 16 of the 18 implants were successful. After spike sorting, manual curation and additional quality control (see Materials and Methods), we obtained 63 units (from p=6 probes) in wS1 and 104 (p=5) in PER. From the same recordings, we also obtained 87 (p=8) units in the secondary somatosensory cortex (S2), 59 (p=5) in temporal association area (TEa), 49 (p=4) in primary auditory cortex (AUDp), 58 (p=5) in dorsal auditory cortex (AUDd), and 73 (p=5) in ventral auditory cortex (AUDv).

### Single-cells exhibit trial window and stimulus-specific responses

Given the repetitive nature of the stimulations, we wanted to assess how responses would vary across the 5s stimulation window and between the stimulation types. Figure 2 shows the firing patterns of three neurons in wS1 and PER, exemplifying the diversity of the observed responses in the two areas. Some neurons in wS1 responded consistently to the repeated whisker deflections but exhibited distinct responses for ipsilateral and contralateral whisker stimulation (e.g., see Figure 2A and B). In other units, the response evoked by the first pipette movement was instead different from that elicited by the subsequent movements (e.g., see Figure 2C). The latter type of response was found more frequently in PER neurons, which typically only responded to the first whisker deflection in the sequence (see Figure 2E and D). By contrast, consistent responses to every whisker deflection were rarely observed (e.g., see Figure 2D). Furthermore, PER responses were generally similar between trial types (i.e., for the two stimulation sides). Overall, these example response patterns at the single cell level suggest that the stimulus features are encoded differently in the two regions.

**Figure 2:**
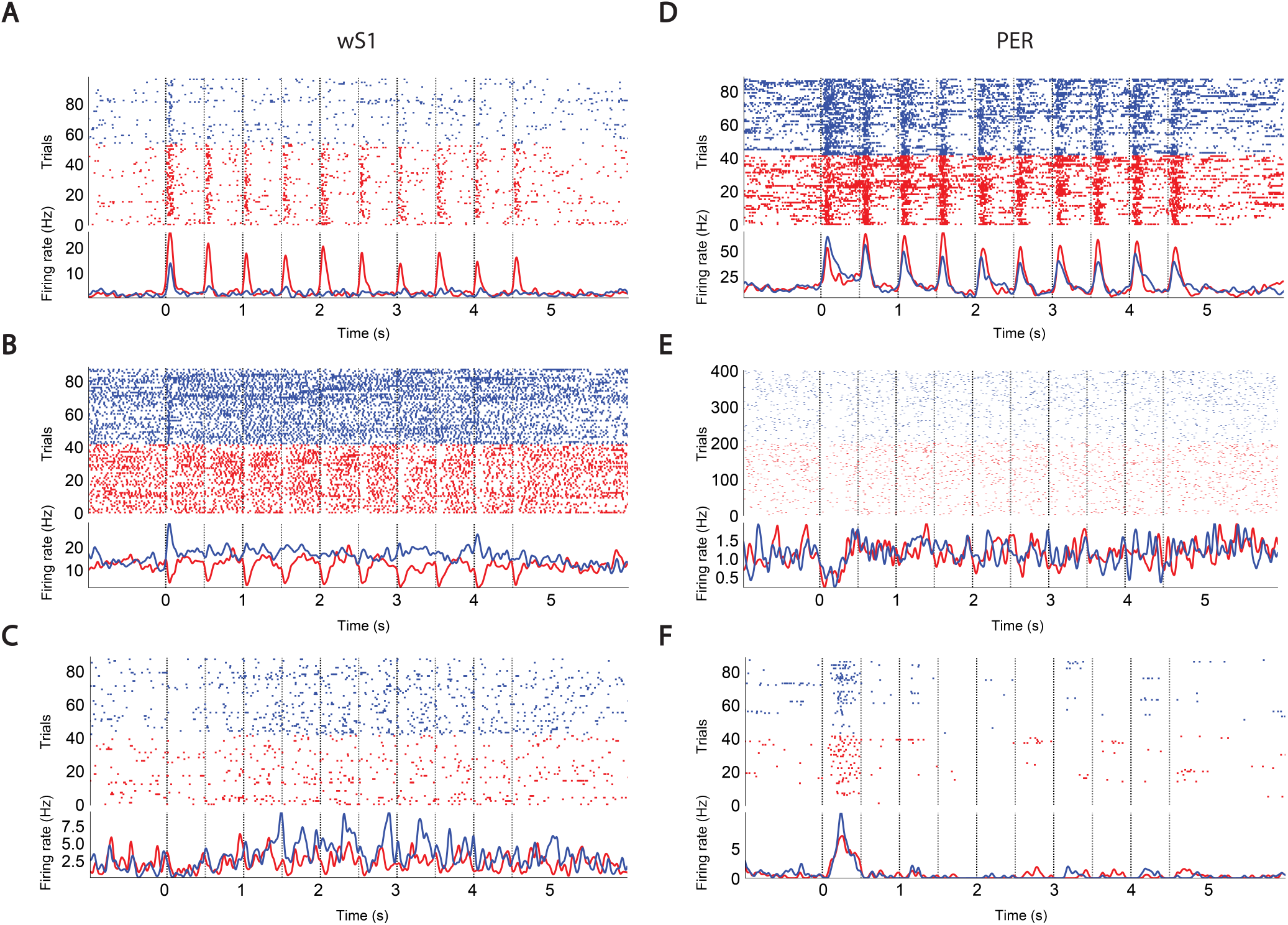
Single-unit responses to tactile stimuli. **A-C:** Raster plots and mean activity traces of three example neurons from wS1. Although the selection of trial type was pseudorandom, they have been separated in the raster plots for visualisation purposes. Contralateral trials and traces are in red and ipsilateral in blue. **D-F:** As **A-C**, but for three PER units.

**Figure 3:**
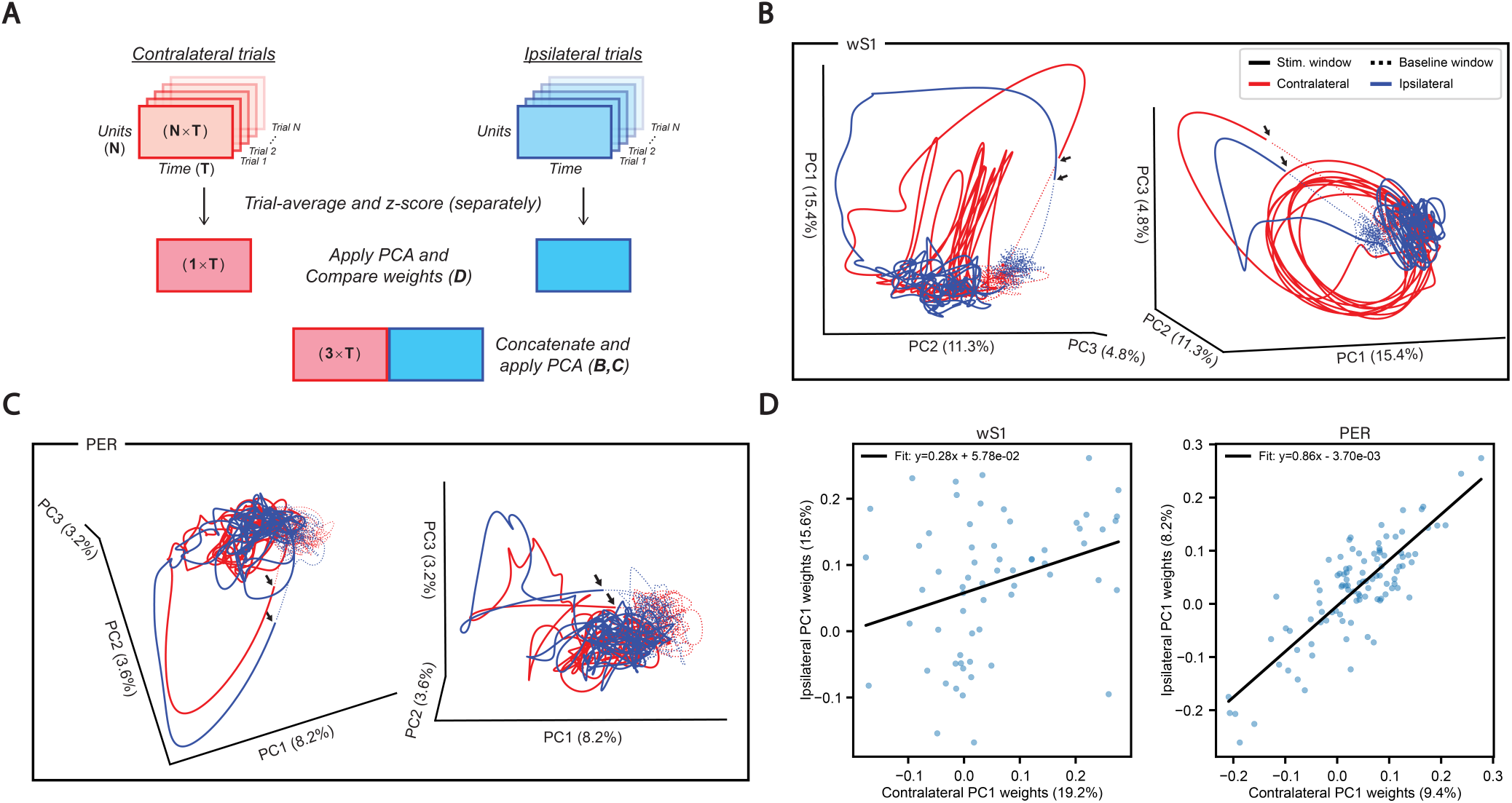
PCA decomposition of the population responses during a window of [−2.5–7.5]s relative to stimulus onset. **A**: Schematized explanation of the methodology to produce panels. **B**-**D** for each region. The dimensions of the output data in each step are indicated for the contralateral trial data and is identical for ipsilateral trial data. Trial-averaged data for each stimulus type were smoothed and z-scored (see Materials and Methods) before applying PCA either individually or upon concatenating the data first. **B**: Two views of a PCA projection of the concatenated, trial-averaged data in wS1 for the first three PCs. Solid lines correspond to the responses during the 5s stimulation window while dashed lines correspond to the baseline. The contralateral case is in red and ipsilateral in blue. Black arrows indicate the onset of the first stimulation, slightly offset due to smoothing artifacts. Percentages within axes labels denote the explained variance of each component. **C**: Same as in **B**, but for PER responses. **D**: Scatter plots comparing the weights from the first principal component (PC) on applying PCA separately across trial types. Each point (blue) represents a neuron’s contribution to the PCs, with a regression line (black) summarizing this across the population. Axis labels also denote in brackets the percentage of explained variance for the corresponding principal component.

### Population dynamics are qualitatively distinct in wS1, but not in PER

Next, we used PCA to examine the population-level dynamics produced by the whisker stimulation sequence in the neuronal representational space. For each stimulus type (contra- or ipsi-lateral), we z-scored the trial-averaged activity of each neuron, creating a matrix of neuronal population vectors across 1ms time bins (Figure 3A). For each region, the PCA analysis was performed in two different ways: 1) the two matrices were concatenated prior to using PCA; and 2) the two resulting matrices were separately decomposed using PCA.

The former approach was used to visualise the population response dynamics for the two stimulation sides. More specifically, we concatenated the corresponding matrices time-wise, we applied PCA across the neuronal axis, and we plotted the projections of the population vectors during the stimulation epoch in the space defined by the first three principal components (Figure 3B,C). Overall, wS1 dynamics were distinct between stimulation types, whereas they appeared largely overlapping in PER. For both regions and stimulation types, the onset of the stimulation window produced the largest deviation from baseline activity (Figure 3B,C; black arrows). Moreover, subsequent stimulations evoked distinct response patterns between the stimulation sides (i.e., ipsi- vs contralateral trials) in wS1, a feature not observed in PER.

To explore this further, we applied PCA separately to the two matrices to determine whether the neuronal populations were similarly activated by the two stimulation types. Specifically, we extracted the weights of each neuron from the first principal component of the two PCA decompositions (Figure 3D). For wS1 neurons, there was only a weak, although significant, correlation (*R* = 0.31*, P* = 1.34 × 10*^−^*^2^), while in PER the correlation was much stronger (*R* = 0.83*, P* = 7.9 × 10*^−^*^28^). This indicates that the stimulus representation in PER was way more stable across stimulation sides as compared to the wS1 representation.

Together, these observations suggest that wS1 and PER respond differently to the same stimulus at the population level, and that the response dynamics in PER are more similar across different stimulus conditions (i.e., stimulation sides) than in wS1.

### Cross-conditional stimulus-baseline decoding supports the notion of side invariance in PER representations

The PCA analyses shown in Figure 3 suggests that PER neurons encode the presence of tactile stimuli in a way that is invariant with respect to the stimulation side. However, this conclusion is based on visual inspection of the population response trajectories in the low dimensional space defined by a small set of principal components. To better quantify this phenomenon, we applied classifiers to discriminate the stimulus epoch from the baseline epoch (i.e., intertrial intervals between stimulation epochs) based on the responses of the recorded neuronal population in a cross-conditional manner. That is, we assessed how the decoding accuracy was affected when the classifiers were trained with neuronal population vectors produced by stimulating one side of the face (e.g., ipsilateral) and tested with population vectors elicited by stimulation of the other side (e.g., contralateral). These performances were compared to those obtained when the classifiers were trained and tested with population vectors obtained by stimulation on the same side of the face (within-condition decoding). The rationale was that, if performance remained similar for the cross-conditional and within-condition decoding, this would indicate that population representations were similar between stimulation sides.

To do this, we trained multiple SVM classifiers to distinguish stimulus and baseline firing patterns along the stimulation window (Figure 4A). First, *N* = 30 units were randomly selected and trials were randomly partitioned into two groups. The activity in each group was then trial-averaged, producing two 2D matrices of 5ms-binned firing rates (see Materials and Methods). For each group, an SVM was trained on the first 150ms of data (30 bins) following stimulus presentation along with 150ms of additional data, taken starting at a random point within the baseline window. The classifier was trained to discriminate the population vectors belonging to the stimulus epoch from those belonging to the baseline epoch. The corresponding data of the other group was used as the test set to assess the classifier performance. This same approach was applied across the entire stimulus window, with a step size of 50ms. We performed this procedure 100 times, sampling, for each run, a new subpopulation of 30 neurons and a new trial partition. We then averaged the resulting 100 accuracy values to obtain our final measure of discrimination accuracy between baseline and stimulus epochs. As mentioned above, the train and test population vectors could either be produced by the stimulation being applied to the same side (within-condition decoding) or to different sides (cross-conditional decoding).

Figure 4B illustrates the performance of within-condition and cross-conditional decoding when using wS1 neurons. In both plots, the decoding accuracy obtained by training and testing on data from the same stimulation side (coloured curves) was different from that obtained when training and testing data were from different stimulation sides (black curves). In particular, the cross-conditional accuracies were substantially lower than the within-condition accuracies, especially after the first whisker deflection. This is consistent with the single-unit responses (Figure 2) and the population response trajectories (Figure 3) observed in wS1. Figure 4D quantifies the difference in the average decoding performance over time when train and test data belonged to the same stimulation side vs when they belonged to different stimulation sides (e.g., the red curve in D is the difference between the black and red curves in the left plot of B). These differences are largest at the onset of each piezo movement and reduce greatly thereafter, in line with the dynamics observed in the PCA trajectories (Figure 3C). Overall, the mean difference over time between cross-conditional and within-condition accuracies was large and significantly lower than zero (p=0.001; permutation test; see Materials and Methods). This indicates that the information encoded by the wS1 representation during contralateral stimulation differed substantially from that encoded during ipsilateral stimulation, and vice versa.

In contrast, when using PER neurons (Figure 4C), the decoding performance was more similar across pairs of train and test data, i.e., no matter whether train and test population vectors belonged to the same (coloured curves) or to different stimulation sides (black curves). Moreover, no clear pattern emerged from the differences in decoding performance over time (Figure 4E). The distributions of the accuracy differences clustered more tightly around 0%, as compared to the case of wS1. When the test side was the ipsilateral (blue curve in Figure 4E), the mean accuracy difference over time was not significantly larger than zero (p=0.2547; permutation test). When the test side was the contralateral one (red curve), the mean accuracy difference was slightly but significantly negative (p=0.001; permutation test). Yet, it was one order of magnitude lower than what observed in wS1 (compare to Figure 4D; note the different scale on the ordinate axis). Overall, this indicates that, differently from wS1, PER encoded the presence of the stimulation sequence in a way that was very similar (i.e., invariant) with respect to the stimulation side.

### PER side invariance illustrated through stimulus type decoding

Here, we assess response similarity more directly by training SVM classifiers to decode the side of the face being stimulated. The approach is similar to the one described before, except that the splitting of trials into two groups is done for both trial types together. Essentially, the other trial type takes the place of the baseline data in the previous approach.

Figure 5A shows the decoding accuracy along the stimulus window. In the case of wS1, decoding accuracy was high, peaking shortly after each stimulation and then decaying to chance level within about 200 ms. By contrast, decoding accuracy remained close to chance level for PER along the whole stimulation epoch. To assess the statistical significance of these trends, we compared these accuracies to empirical distributions of chance performances obtained by shuffling the labels of the stimulation side (permutation test; see Materials and Methods). This analysis showed that decoding accuracy was not different from chance throughout the entire stimulation window for PER. Instead, stimulation side could be decoded significantly above chance throughout most of the stimulation period based on wS1 activity.

**Figure 4:**
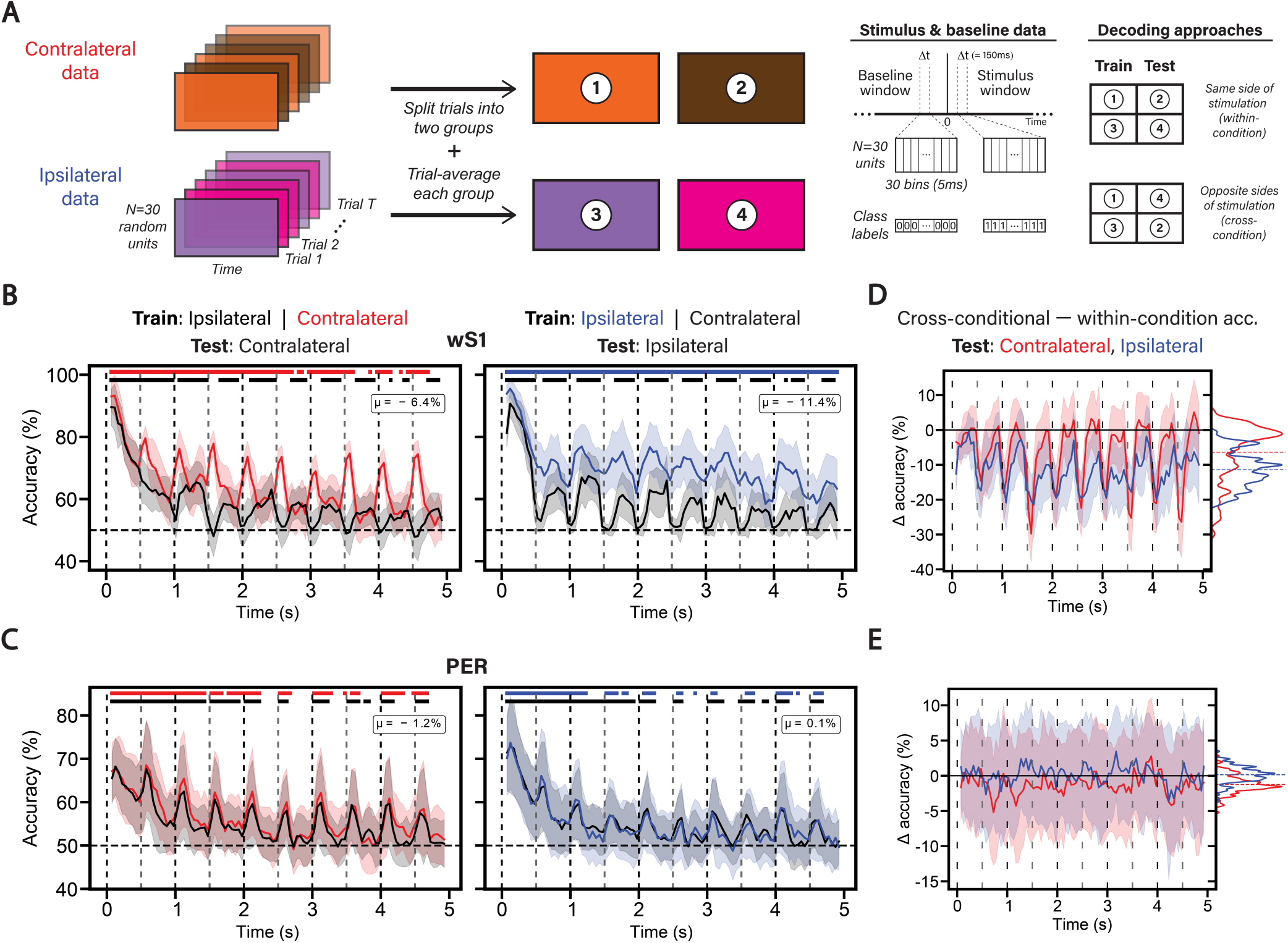
Cross-conditional decoding analyses of stimulus and baseline windows. **A**: Schematic illustrating the preprocessing, stimulus and baseline data extraction, and the within- and cross-conditional approaches for evaluating classification accuracy (see Materials and Methods for details). On the right, the splitting of trial groups into train and test sets is also repeated with the opposite assignment of train and test sets. **B**: Two plots corresponding to the possible combinations of trial type (i.e., ipsi- and contralateral) data chosen for training and testing in wS1. Each plot contains two curves corresponding to the average decoding accuracy (obtained over 100 independent runs of training and testing a classifier, each with independent choices of neurons and trial splitting) over the 5s stimulation window using a 150ms sliding window with a 50ms step size. The coloured curves correspond to the average decoding accuracy obtained from training and testing on data from the same trial type, while the black curves correspond to training on the opposite trial type. In each 150ms window, the corresponding 5ms-binned stimulus data, and randomly chosen baseline data, are used for training and testing each classifier (see Materials and Methods). Shaded regions around the average accuracies signify the standard deviation over the 100 independent classifiers. Vertical dashed lines are in black and grey, and mark the two directions of stimulation. Horizontal dashed line marks the theoretical chance level, with solid bars above the plot indicating regions where accuracy was significantly above a corresponding empirical null distribution obtained through shuffling class labels. Each plot contains text indicating the average difference between the cross-conditional and within-condition decoding accuracies. **C**: As **B**, but for PER. **D**: Two curves showing the difference in average decoding accuracy over time between the cross-conditional and within-condition cases, coloured by the trial type data used for testing the classifiers, with the average difference over time indicated by a dashed line. On the right of each panel is a density plot of these differences, with the average difference over time indicated with a dashed line. **E**: As **D**, but for PER.

**Figure 5:**
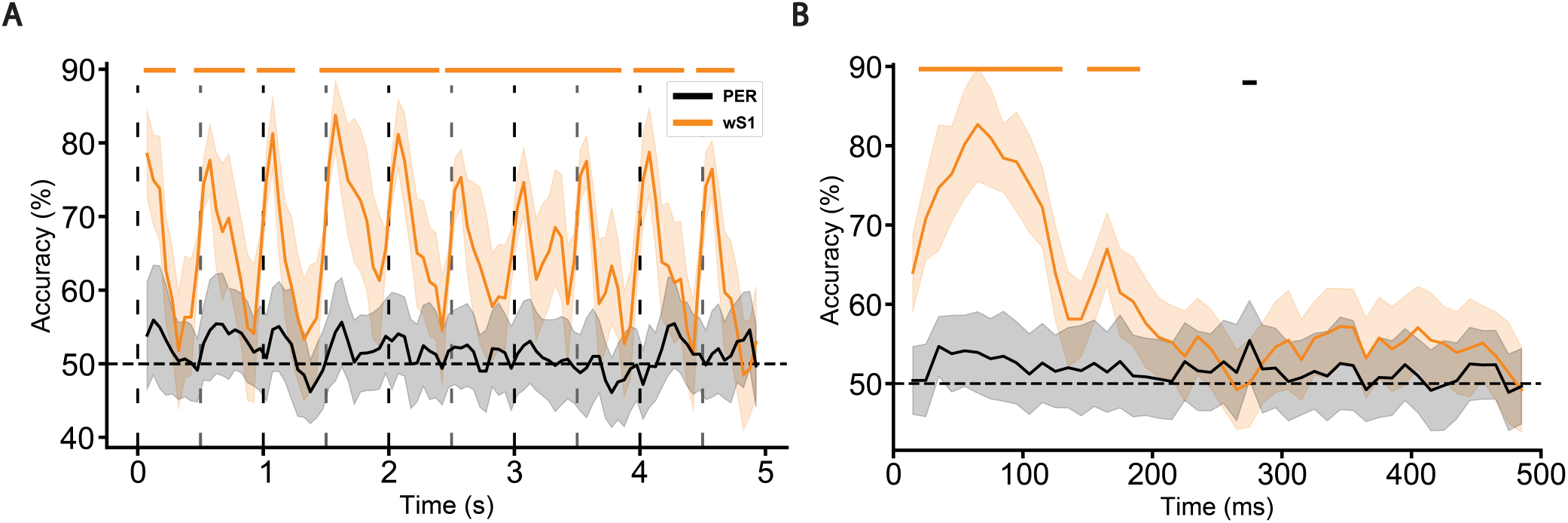
Assessing the ability of decoders in classifying each of the two stimulus features. **A**: Average decoding performance for classifying the side of stimulation in wS1 (orange) and PER (black). The vertical dashed lines mark the times of stimulation, with the two directions of motion indicated by the colour of the lines. The horizontal dashed line corresponds to the theoretical chance level. The solid lines above the plot mark the temporal regions where decoding accuracy was above an empirical chance level obtained through shuffling class labels (see Materials and Methods) and these lines are colour-matched to each region. **B**: Average decoding performance for classifying the direction of stimulation in contralateral trials. This figure follows the same conventions as **A**, but is limited to a 500ms window as the decoding accuracy was relative to movement onset for each of the 10 movements in the stimulation window.

Importantly, wS1 sends direct projections to PER and also lies at an earlier stage of processing in the somatosensory hierarchy. As such, it is interesting that, despite there being sufficient information in wS1 neurons to reliably decode the side of the face being stimulated, such information is virtually absent in PER. This observation is reminiscent of similar findings in the visual domain, where luminance and contrast information have been shown to decrease along a the rat homologue of the ventral visual stream, from the primary visual cortex to extrastriate area LL - a trend that has also been found in convolutional neural networks (Tafazoli et al., 2017; Muratore et al., 2022). In this regard, our results show that, along the somatosensory processing hierarchy, information about the side of the face being stimulated is absent at the stage of PER, meaning that the region represents the stimulus in a side invariant manner.

### Extending invariance observations beyond side through stimulus direction decoding

Next, we tested whether responses in PER show invariance to other physical features of the stimulus. In our experimental design, an object moves along the rostro-caudal axis which allows to observe neuronal responses to movements in two directions along this axis. We therefore used the same neurons to assess performance in decoding stimulation direction. Since both directions of movement are represented in each trial, we made some changes to the decoding approach (see Materials and Methods). Briefly, we split the 5s stimulation window into ten 500ms chunks, where each chunk is aligned to a stimulus movement direction, either forward or backward. Decoding accuracy was therefore calculated relative to movement onset, between 0 and 500ms. As this approach provides five times more data compared to stimulation side decoding, we used 30ms windows instead of 150ms, with a step size of 10ms instead of 50ms. Moreover, since the key feature is direction, we only used trials involving contralateral stimulations.

The result of this analysis is shown in Figure 5B. In the case of PER, decoding accuracies were around chance level, with only one bin, centered at 275ms after stimulus onset, yielding an accuracy that was significantly above the shuffled distribution. By contrast, the decoding accuracy afforded by the wS1 representation was much larger, featuring two peaks where it was significantly above chance, during the initial epoch of the response following the whisker deflection. Information about stimulus direction is therefore present in wS1 but absent in PER, thus indicating that the latter is invariant to this stimulus feature too.

### Invariance in PER is neither inherited nor a universal property of higher-order processing

That stimulation side and direction cannot be reliably decoded in PER does indeed indicate invariance to those features. Somatosensory information is transmitted from wS1 to S2 and TEa, which provide prominent inputs to PER (Burwell and Amaral, 1998). Our Neuropixel recordings granted us access to all of those areas, allowing us to test whether response invariance in PER is inherited from upstream regions and where it emerges along the hierarchy.

To do this, we applied the same decoding analyses for side and direction in both S2 and TEa. The results of these analyses show that both stimulus features are reliably decodable from the S2 population activity (Figure 6), particularly stimulus direction, which was decodable for almost the entire 500ms window following wisher deflection. Also for TEa, although accuracy was generally lower than in S2, there were multiple bins significantly above chance, particularly in decoding stimulation side.

**Figure 6:**
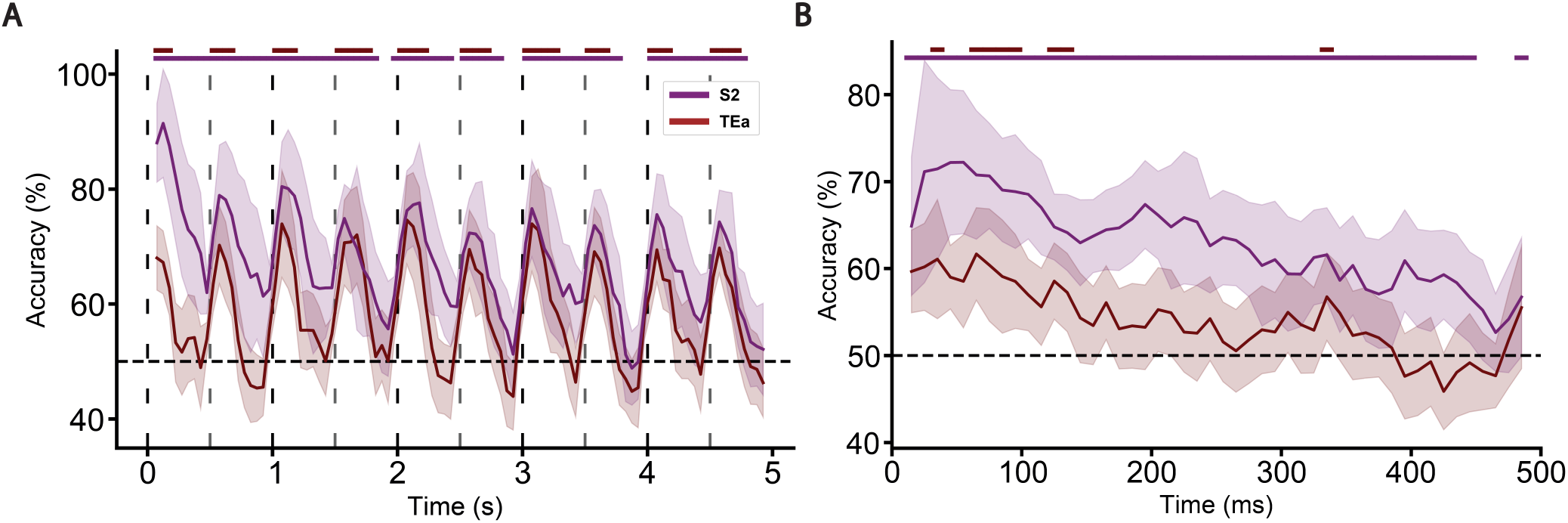
Assessing stimulus side (**A**) and direction (**B**) decodability using neurons in S2 (purple) and TEa (maroon). See Figure 5 and Materials and Methods for details.

The percentage of time bins that were decodable above chance for both stimulus features, based on the population activity recorded from all four regions are summarised in Figure 7. For the decoding of the stimulation side, S2 had a percentage of significant bins that was similar to wS1. Strikingly, S2 had instead a much higher percentage for direction decoding. For both features, decoding accuracy decreased in TEa, although it remained significantly above chance level, suggesting that fully invariant responses emerge in later stages of the cortical hierarchy (Figure 6). Such a level of invariance was indeed observed in PER, as demonstrated by our decoding analysis shown above.

**Figure 7:**
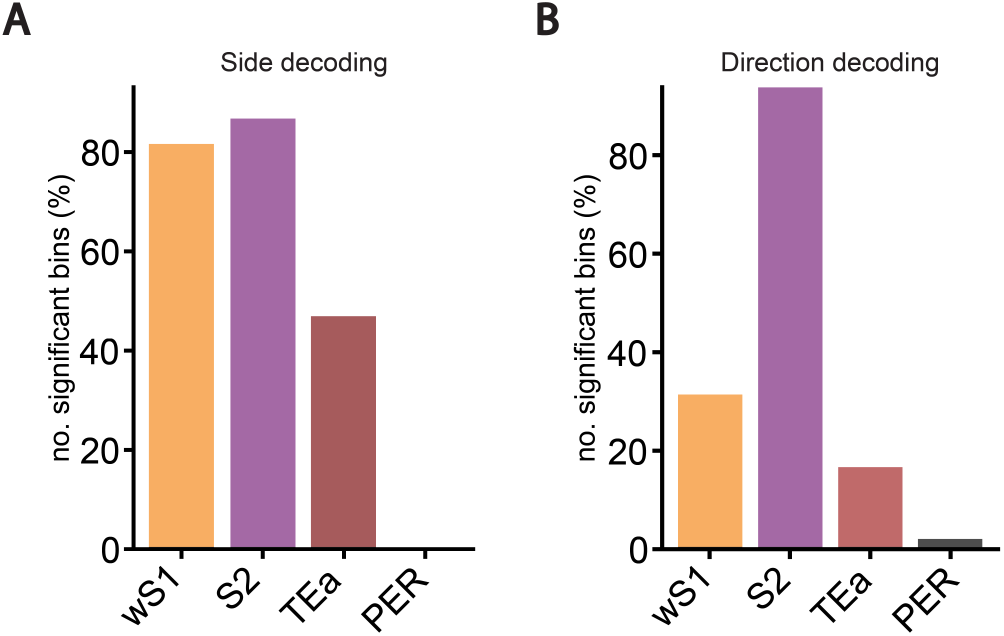
Bar charts summarising the results of the stimulus feature decoding analyses (Figures 5 and 6). They show the percentage of temporal windows whose average accuracy were significantly above chance in all four regions. **A**: Results for decoding the side of the face being stimulated. **B**: Results for decoding the direction of movement along the rostro-caudal axis for contralateral trials.

It should also be considered that the movement of the pipettes also produces an audible click, which differs in sound between pipettes and also based on their direction of movement. To check whether these clicks evoked responses that were also informative about the stimulus features (i.e., stimulation side and direction), we applied all the decoding procedures to neurons that were also recorded in primary (AUDp) and secondary (AUDd and AUDv) auditory cortices (Figure 8). These analyses showed reliable decoding of both stimulation side and direction. Together, these findings strengthen the claims that fully invariant representations of stimulus presence emerge in PER, suggesting an important role for PER in encoding the awareness of perceptual objects (Burke et al., 2012).

**Figure 8:**
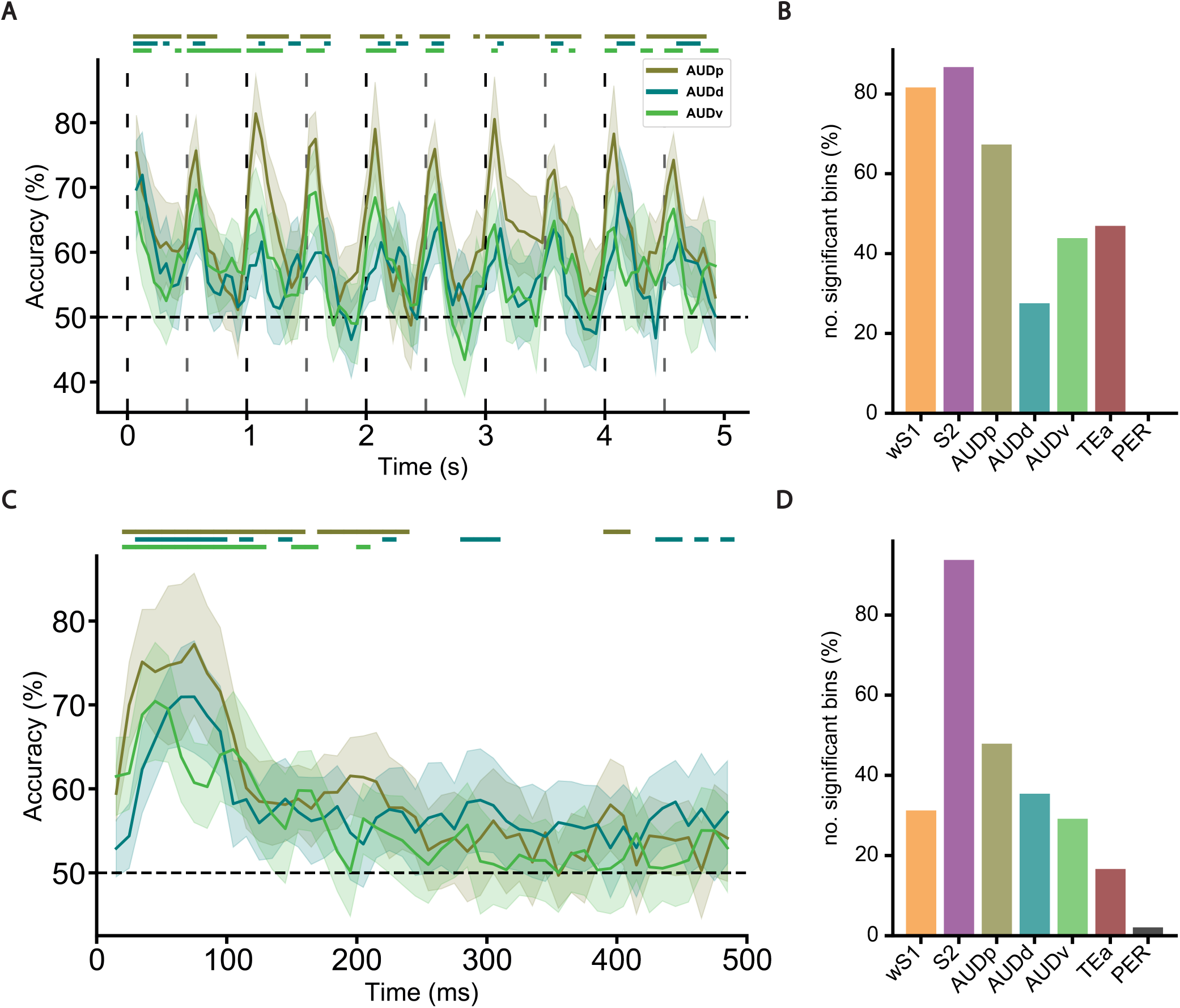
Applying the same stimulus feature decoding approaches in AUDp, AUDd, and AUDv. **A**: Average decoding accuracy for decoding the side of stimulation, coloured by region. Dashed horizontal line is the theoretical chance level and vertical dashed lines indicate the times where stimulations occurred, which is in black and grey to label the two stimulus movement directions. Solid lines above the figure show the times where decoding accuracies were above their corresponding empirical chance distribution obtained through shuffling class labels. Error bars around average accuracy traces are the standard deviation. **B**: Bar plot showing the percentage of decoding time windows whose average accuracy was significantly above chance for both the auditory regions and all other recorded regions. **C** and **D**: As (**A**) and (**B**), respectively, but for decoding the direction of stimulation in contralateral trials.

### Abstraction of stimulus features revealed through clustering of neural response patterns in PER

The previous sections established that PER is the first region in the somatosensory processing hierarchy where invariance to the two features of our stimuli emerges. To expand our account of perceptual processing in PER, we also sought to uncover the neural response dynamics that underlie this phenomenon. Although we have an account of this at the population level through PCA (Figure 3D), the low fraction of variance explained by the first principal components points to a rich and diverse response dynamics. Therefore, we applied clustering analysis to identify distinct, characteristic classes of response dynamics shared amongst neurons. The number of such classes and the shape of their firing patterns provided information on the extent to which the response patterns were diverse, but also served as a condensed account of the single-cell response patterns. Accordingly, this approach provided a reasonable trade-off between pure single-cell analyses and population-level accounts.

In our analysis, we used K-means to cluster neurons sharing similar response patterns. First, baseline-subtracted smoothed firing rates were computed and z-scored over a window spanning −1s to 6s from stimulus onset (Figure 9A). For each trial type, the number of temporal dimensions was reduced using PCA to extract the first three principal components (see Materials and Methods). Accordingly, the size of the data was reduced from *n* × 7000 to *n* × 3, where *n* is the number of units in PER. It should be noted that this application of the PCA is different from the one used in the analysis of Figure 3. In that case, PCA was applied along the neural dimension, while here it is applied along the temporal dimension. This allowed projecting all the recorded units in each area in a low-dimensional space defined by temporal features of the tactile stimulation sequence. These reduced outputs were combined across stimulation types and processed through K-means. Using silhouette analysis and elbow plots, *K* = 4 clusters was determined as the most appropriate choice.

**Figure 9:**
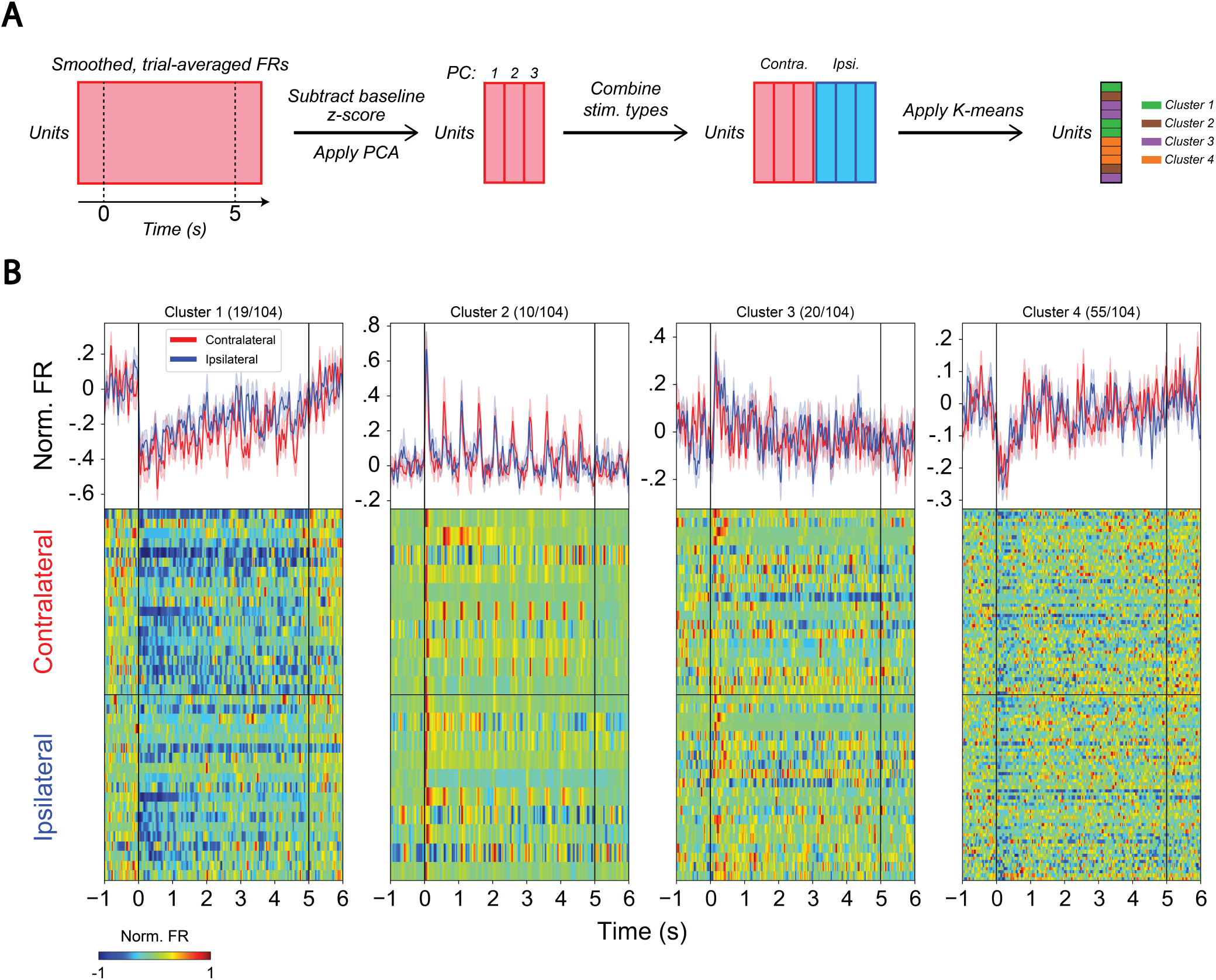
K-means clustering of neurons in PER using their neural response patterns. **A**: Schematic outlining the procedure. Briefly, the firing rates of trial-averaged, smoothed, and baseline-subtracted neurons were z-scored. PCA was then applied to the time dimension. This was done separately for each trial type and the final output was used to cluster neurons using K-means. **B**: Four sets of plots for each of the resulting clusters. In each set, the bottom two panels show a heatmap of trial-averaged, smoothed, and baseline-subtracted neuronal firing rates that were normalised according to their absolute maximum firing rate. The neurons in each panel are row-matched for comparison. The top panel shows the neuron-averaged firing rate per trial type, with shaded regions indicating the standard error of the mean. Above each panel is the cluster label along with the fraction of units in each cluster.

The resulting clusters are shown in Figure 9B for PER. As expected, the cluster-averaged response patterns are similar between the trial types (i.e., contra- and ipsilateral) in all four clusters. Interestingly, only one cluster, containing just under 10% of the population, showed clear responses to all whisker deflections (Figure 9B; Cluster 2). In the case of the first cluster (also including a small subset of cells), the response dynamics featured a slow recovering back to baseline activity after an initial decrease following the first tactile stimulus in the sequence. These two clusters differed from the latter two (Figure 9B; Clusters 3 & 4), which together comprise around 70% of the population and whose responses were instead transient, following the first whisker deflection.

Intriguingly, the firing rate dynamics of the first two clusters are similar in shape to the two primary features of the decoding accuracy profile for the stimulus vs. baseline analysis (Figure 4C), namely the decay envelope and the peaks at each stimulation time. To determine whether these subpopulations are responsible for these features, we repeated the stimulus-baseline decoding analysis but removed these neurons from the sampling pool (Figure 10, black curves). For both trial types (i.e., contralateral and ipsilateral), decoding accuracy was above chance after stimulus onset but quickly returned to chance levels and remained there for most of the stimulus window duration—very different dynamics, compared to the one found using the whole neuronal population (coloured curves). Thus, our decoding strategy suggests that only the first two clusters (despite the scant number of units) provide information about the duration and dynamics of the stimulus, while the latter two clusters provide information about the onset of the stimulation period.

**Figure 10:**
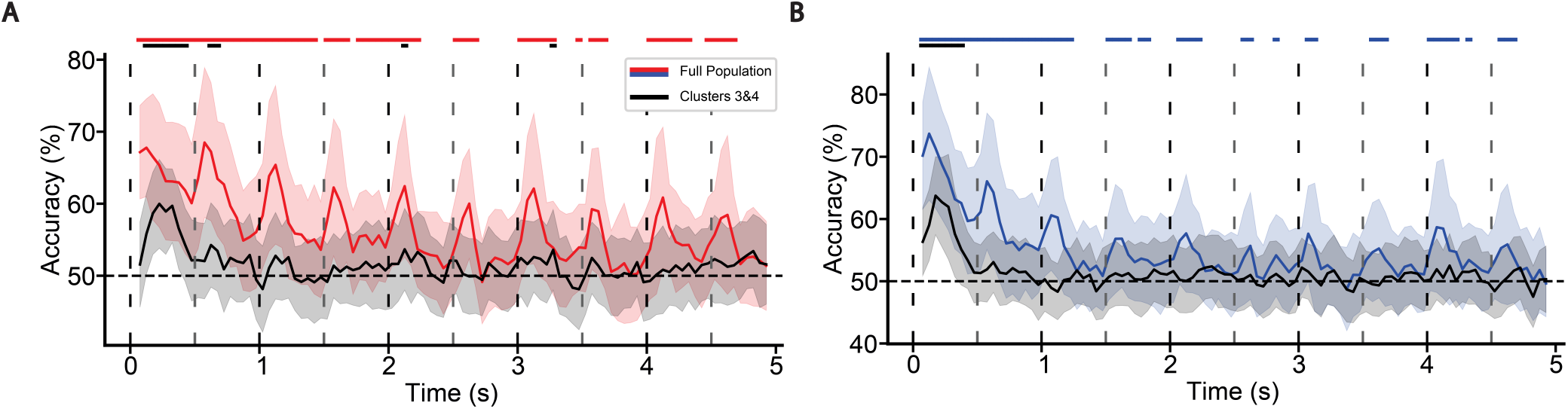
Comparing stimulus-baseline decoding accuracy between the full population and a subpopulation of neurons in PER (Figure 9; Clusters 3 and 4). **A**: Average decoding accuracy using contralateral trials for the full population (red) and subpopulation (black). Error bars denoting the standard deviation. Vertical lines indicate the times where the stimulations occurred, coloured by the direction of movement. Horizontal dashed line is the theoretical chance level. Solid bars above the plot denote each time period where decoding accuracy was above the corresponding two-sided 95% confidence interval obtained through shuffling. These confidence intervals were maxT adjusted to control for the family-wise error rate. The bars are coloured to match the data they refer to. **B**: As (**A**), but for ipsilateral trials. Here, the full population is coloured in blue.

## Discussion

A longstanding topic of debate within the PER literature is the extent to which its role is perceptual or mnemonic (Eacott, Gaffan, et al., 1994; Buffalo et al., 1998; E A Murray and T J Bussey, 1999; Stark and Squire, 2000; M J Buckley et al., 2001; T J Bussey et al., 2005; Hampton, 2005; Levy et al., 2005; Squire et al., 2006; T Bussey and L Saksida, 2007; E A Murray, T J Bussey and L M Saksida, 2007; Suzuki, 2009; Graham et al., 2010; Knutson et al., 2012; Ahn and I Lee, 2017; Bonnen et al., 2021). In this literature, however, virtually all of the works that performed electrophysiological recordings also incorporated task components that warrant mnemonic processing (see (Burke et al., 2012; Thome et al., 2012; Baruchin et al., 2018) for examples to the contrary). From this standpoint, our recordings provide a valuable contribution to the debate because they were conducted in mice engaged in a passive sensory stimulation paradigm. This enables a more refined investigation into the perceptual component of PER activity as mnemonic processing is not instrumental in our experiment. Using this approach, we found that PER neural responses to the stimulus are invariant to its movement direction or the side of the face being stimulated. Along the somatosensory hierarchy, we also showed that these invariances first emerged in PER. Moreover, clustering analysis of PER response patterns revealed that over 90% of the neurons were contained in clusters whose response dynamics were not perturbed by the additional stimulus movements after stimulation onset. Further, the majority of this subpopulation responded only to the onset of the stimulus sequence, with the rest encoding the presence of the stimulation epoch. Since these responses are invariant to the basic stimulus features, it means that information about the particular side of stimulation and direction is abstracted away in favour of encoding the broader temporal structure of the stimulation protocol itself, i.e., the transition from the baseline period with immobile stimuli to the stimulation window where one of the stimuli are in motion. Interestingly, these findings are consistent with those of a recent study reporting that a consistent fraction of PER neurons encodes the presence of 3d objects in a circular track in a way that is invariant with respect to object identity (Burke et al., 2012).

Along the somatosensory hierarchy, invariant responses become more prominent, in part because of the coarser topographic representation leading to broader receptive fields of neurons in higher order cortices (Carvell and Simons, 1986; Kwegyir-Afful and Keller, 2004; Ashaber et al., 2014; Goldin et al., 2018; Hubatz et al., 2020; Parra et al., 2022). In line with these observations, anterograde tracing experiments from wS1 showed that it projects to a narrow band located rostrally in PER (Aronoff et al., 2010). It is therefore likely that PER has no topographic representation of the whisker pad, and this architectural feature could contribute to the invariance seen in our study. Another architectural feature that seems relevant to the interpretation of our results is the strong interhemispheric connectivity of PER with somatosensory areas of the contralateral hemisphere (Fenlon et al., 2017). Indeed, interhemispheric connectivity has been shown to underlie invariant responses in row A of wS1 (Montanari et al., 2023). PER shows similar connectivity patterns with other sensory modalities (Burwell and Amaral, 1998), suggesting that similar invariant responses might be found in other modalities.

Invariant responses are a prerequisite of object perception and recognition (DiCarlo, Zoccolan, et al., 2012). Lesion studies have shown that PER is involved in these processes (E Murray and Mishkin, 1986; Meunier et al., 1993; Eacott, Gaffan, et al., 1994; T J Bussey et al., 2002; T J Bussey et al., 2003; Bartko et al., 2007a; Bartko et al., 2007b) and our results might thus provide important insights into the mechanisms by which PER participates in them. However, it was recently shown that PER does not encode object-specific information in rats performing a multisensory object-recognition task (Fiorilli, Marchesi, et al., 2024). Although seemingly contradictory to our findings, the rats in their experiments were highly familiar with the objects, whereas our mice had never experienced the stimulation paradigm before. This difference in stimulus novelty is a key factor, as the effects of stimulus preexposure on PER activity were inherently absent in our case. A notable example concerns the feedback projections from the deep layers of the PER to layer 1 in the primary somatosensory cortex, which have been shown to facilitate learning in a microstimulation task in mice (Doron et al., 2020). Chemogenetic inhibition of these projections did not significantly affect the performance of expert mice, which are habituated to the stimulus, but had profound effects on learning in näıve mice.

Alternatively, PER might contribute to building a sensory context onto which to map task-relevant features such as reward. In this scenario the movement of an identical object in the two sides of the face is represented equally unless a valence is assigned to one stimulus. Indeed, Fiorilli et al. (2024) found that PER populations encode choice and reward (Fiorilli, Marchesi, et al., 2024). These findings are supported by anatomical data showing a strong reciprocal connectivity of PER with the amygdala, whose inputs might be important to build sensory-reward associations (Burwell and Amaral, 1998; Pitkanen et al., 2000; Agster and Burwell, 2009). Neurophysiological experiments in vitro and in vivo demonstrated that amygdalar inputs are necessary to propagate sensory information towards the LEC and the memory system (Kajiwara et al., 2003; Paz et al., 2006). Our stimulus-baseline decoding shows that the low decoding accuracy of input identity is not due to a general absence of stimulus representation. These results suggest that PER can represent a sensory context independently of the physical features of the stimulus.

Beyond reward encoding, the connectivity of PER suggests that multisensory processing might represent another mechanism for object identity or context encoding in PER. Indeed, multisensory processing has been shown to support object recognition in PER (Winters and Reid, 2010; Jacklin et al., 2016). All methods of tactile stimulation produce sound, and both modalities are present in our experimental design as shown by neural responses in wS1 and the three AUD regions. Future experiments focused on different audio-tactile or visuo-tactile responses will clarify the role of multisensory processing in shaping responses in PER.

A large body of literature shows that PER is not necessary for the recognition of simple stimuli (T J Bussey et al., 2002; T J Bussey et al., 2003; Norman and Eacott, 2005; Bartko et al., 2007a; Ramos, 2016), suggesting little involvement in representation of sensory stimuli at odds with our findings. Supposing that the stimuli in our experiments are indeed not sufficiently complex to necessitate PER utilisation in recognition tasks that would use them, what is the explanation for our observations of invariance and abstracted stimulus representations? One possibility concerns the hypothesised function of unitization performed by PER, i.e., the process of combining multiple sensory features or “items” into the representation of a singular construct (Graf and Schacter, 1989). The features shown to be unitized in PER began with discontinuous auditory segments (Bang and Brown, 2009), and expanded over time to include, for example, paired visual objects (Fujimichi et al., 2010), spatial segments of a task environment (Bos et al., 2017), and word pairs (Haskins et al., 2008). In fact, a recent review proposed that the diverse functions and response patterns in PER can be explained by a consistent unitization operation that is applied similarly to features across domains, therefore allowing PER to be flexibly utilised across various cognitive functions and task demands (Fiorilli, Bos, et al., 2021). Given that we implemented a passive sensory paradigm using simple stimuli—which, as outlined earlier, do not warrant PER utilisation for various cognitive tasks—we believe the observations of side and direction invariance reflect a propensity for PER to unitize stimulus features, in part because the same was not observed in TEa. To explain, consider that our experimental design did not include non-sensory features and that the recordings were conducted in a single session for each mouse. Therefore, it is implausible that the invariant and abstracted representations reflect any mnemonic processing. Rather, these phenomena are likely inherent to sensory processing in PER, presumably by virtue of specialized cell-types (Beggs et al., 2000; Moyer Jr et al., 2002; Courcelles et al., 2024), their microcircuits and the patterns of anatomical connectivity described above. The extent to which these components may contribute should be explored with future modelling studies, with additional experiments assessing whether similar observations emerge when non-sensory features are implemented.

In conclusion, our work demonstrates that stimulus responses are present in PER during a passive tactile stimulation paradigm, that the corresponding stimulus representations are invariant to the two varying stimulus features, and that the majority of neurons cluster into subpopulations whose aggregated response dynamics are indicative of abstracted stimulus representations. We believe that these simple passive sensory investigations are a useful complement to the current electrophysiological investigations that involve more complex tasks or behaviours.

## Conflict of interest statement

The authors declare no competing financial interests.

## Acknowledgments

This work was funded by the European Union’s Horizon 2020 Research and Innovation programme under the Marie Sklodowska-Curie (grants: 885955 and 101063718), and the Research Council of Norway through the Center of Excellence scheme (grant: 332640). We would like to thank Dr. Seán Froudist-Walsh for providing feedback on the manuscript. We would also like to thank Dr. Grethe Olsen for excellent technical support with the histology of the in vivo recordings.

## Notes

### Competing Interest Statement

The authors have declared no competing interest.

